# RNAhub - an automated pipeline to search and align RNA homologs with secondary structure assessment

**DOI:** 10.1101/2025.03.11.642701

**Authors:** Marcin Magnus, William Gao, Nivedita Dutta, Quentin Vicens, Elena Rivas

## Abstract

The complexity in the generation of RNA multiple sequence alignments and assessment of the accuracy of such alignments contributes to the challenges in the utilization of RNA alignments in diverse integrative methods. RNAhub is a freely available user-friendly web server for a reliable generation of RNA multiple sequence alignments and the detection of the presence of structural RNA utilizing evolutionary information. This web-based tool, developed by the integration of existing computational approaches, takes an RNA sequence as input and automatically retrieves and aligns sequences homologous to the input (query) RNA sequence through an iterative and structure-agnostic approach. Based on the alignment, this tool statistically assesses whether the query RNA sequence has a conserved RNA structure using covariation analysis. The web server allows the user to efficiently search the sequence of interest against carefully curated, ready-to-use genomic databases to produce a multiple sequence alignment. Using this alignment, our tool either detects the presence of a conserved structural RNA, finds evidence against the presence of a conserved structure, or cannot make any assessment due to a lack of sequence diversity in the alignment. The web server is freely available at https://rnahub.org.

**Graphical Abstract:** 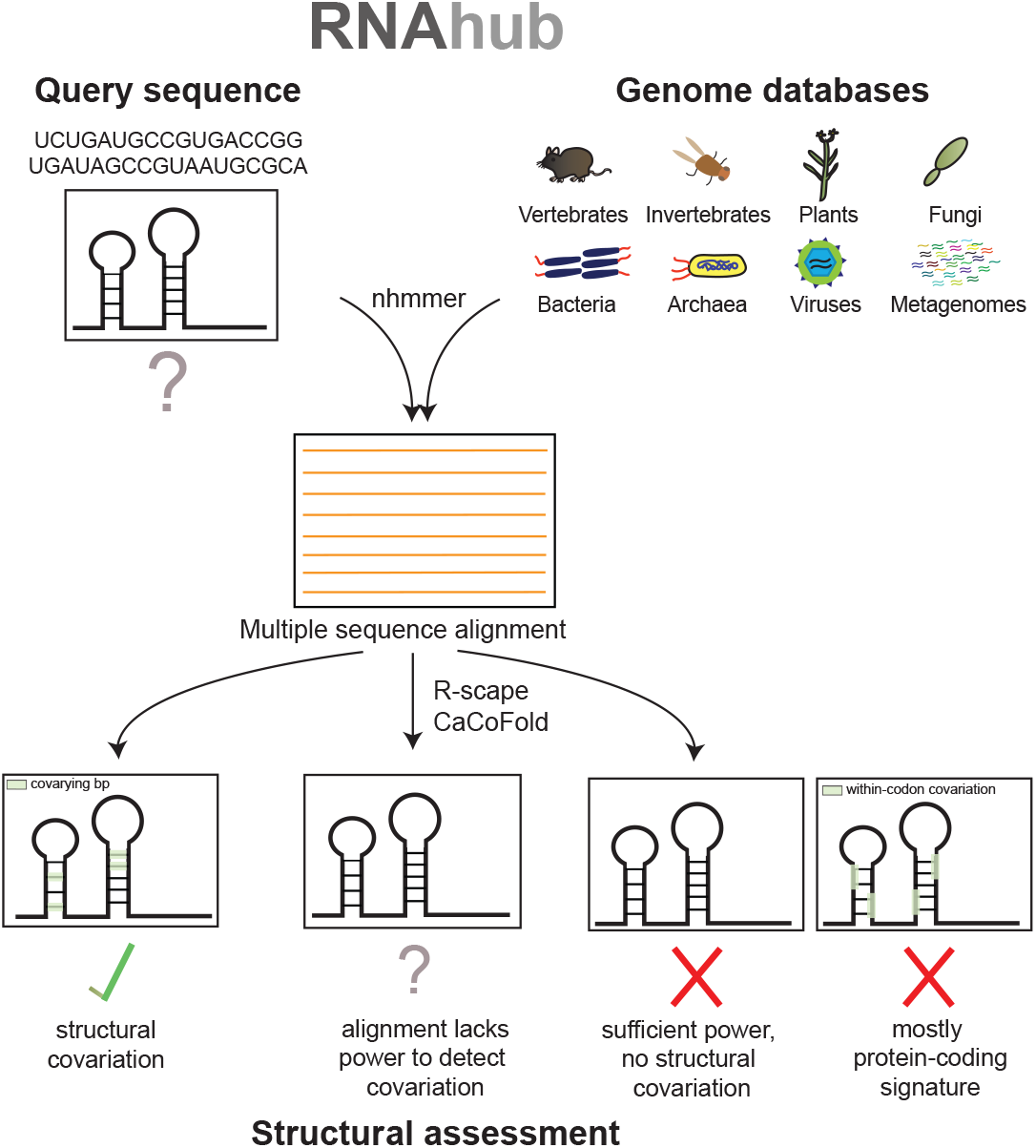

## Introduction

Many non-coding RNAs function through a conserved structure that allows for interactions with other RNAs, proteins, and metabolites. For example, transfer RNAs (tRNAs), which are critical for translation, adopt a cloverleaf secondary structure (Watson-Crick or wobble base pairing) and an L-shaped tertiary structure. This structure allows the tRNA to engage with the protein subunits of the ribosome in a protein-RNA complex [5]. As structure is critical to the function of structural non-coding RNAs, these non-coding RNAs display a distinct evolutionary signature that favors the maintenance of structure via compensatory base pair substitutions [3]. Compensatory base pair substitutions refer to two nucleotide substitutions across a distance of evolutionary time that maintain RNA secondary structure even as the primary sequence changes. For example, in one species, a base pair could be A-U, while in another species, it would be G-C; in either case, the base pairing interaction is maintained. These compensatory base pair substitutions covary (vary with each other) and provide evidence in favor of an evolutionary constraint to maintain RNA secondary structure [16, 22].

Therefore, a critical step in identifying functionally important structural RNAs is to perform a homology search to produce a multiple sequence alignment [1, 9] on which we can identify compensatory base pairs. Accurate RNA multiple sequence alignments are not only crucial for deriving biologically relevant RNA structures but also the key input for the next-generation deep learning methods for RNA structure prediction, such as those inspired by AlphaFold [4].

Generating accurate RNA multiple sequence alignments without prior structural information, as well as evaluating their quality, continues to pose a major challenge. Currently, the identification of novel functional structural RNAs heavily depends on expert knowledge and manual intervention, hindering effective automation and limiting broader adoption by the RNA research community.

Several computational tools, including LocARNA [20] and RNAz [23], attempt to address some of these challenges. LocARNA integrates sequence and structural information to generate RNA alignments by simultaneously aligning sequences and predicting structures. While effective for structured RNA alignments, LocARNA requires initial structural or sequence-based insights to be most accurate. RNAz, on the other hand, predicts functional RNA structures by evaluating alignments based on thermodynamic stability and evolutionary conservation. Despite its utility in RNA structure prediction, RNAz strongly depends on precomputed, high-quality alignments, highlighting the importance of accurate initial sequence alignments. However, none of the aforementioned tools perform homology-based sequence searches; instead, they require an input set consisting of unaligned sequences (LocARNA) or aligned sequences (RNAz).

An online server created for identifying and aligning homologous sequences is rMSA [26]. The server generates multiple sequence alignment searching sequences from the NCBI nucleotide (NT) database and RNAcentral. However, the method uses overly lenient parameters of the structural homology method Infernal [13]. This practice leads to the contamination of alignments with a blend of unrelated sequences from diverse sources such as bacteria, plants, and humans. Moreover, the rMSA server uses predicted structures to construct the alignments that are going to be assessed later for the presence of a structure, which can lead to incorrect statistical assessments and false positives.

To address the mentioned challenges, we introduce a user-friendly web server, RNAhub. The server identifies and visualizes a consensus RNA secondary structure utilizing evolutionary information based on an integration of computational comparative methods involving sequence alignments, analysis of covariation, and detection of the presence of conserved RNA structures. The RNAhub web server builds upon existing tools, such as the structural homology search from Gao *et al*. 2021 [7]. RNAhub builds structure-agnostic alignments using nhmmer [25]. The decision-making process of evaluating the quality of alignments and whether they support the presence of a biologically relevant and conserved RNA structure or not, is performed using R-scape [16]. R-scape is a previously developed computational method designed to identify pairs of positions in the alignment that significantly covary and importantly accounts for covariation due to phylogenetic relatedness in its null model. Rivas *et al*. 2017 [16], demonstrated that R-scape identifies significant covariation in many known structural RNAs but does not detect them in some long non-coding RNAs such as HOTAIR, suggesting these RNAs lack conserved structures. When the tunable expectation value (E-value) is set to a stringent value, such as the default of 0.05, R-scape is very specific and less prone to false positive base pairs than other methods. Later, we demonstrated [7] that R-scape can be used in a yeast genome-wide search of novel conserved structural RNAs with high specificity.

RNAhub implements a workflow in which a query sequence is searched iteratively against a database of choice for homology using nhmmer, resulting in a final alignment that is passed to R-scape to identify significantly covarying base pairs. These significantly covarying base pairs, which have many compensatory base pair substitutions, provide evidence in favor of an RNA whose function is dependent on its structure. The server provides an intuitive interface to carry out all the computational steps mentioned above and offers ready-to-use genome databases for conducting homology searches with the query sequence.

## Materials and Methods

### The overview of the RNAhub workflow

For a query DNA/RNA sequence of interest, the server searches for homology to Rfam [14], a database of conserved structural RNAs. If no Rfam homology is detected, the server conducts iterative homology searches using the structure-agnostic method nhmmer [25] against a broad spectrum of curated genome databases. The resulting alignment is analyzed for evidence of a conserved RNA structure by measuring pairwise covariation (compensatory base pair substitutions) with R-scape [16]. The outcome of the analysis is one of the following (see Figure 1): (i) The RNA is a known structural RNA with homology in the Rfam database. Otherwise, a nhmmer alignment is constructed. By examining the nhmmer alignment with R-scape several outcomes are possible: (ii) The alignment shows covariation consistent with an RNA secondary structure, and a structure is calculated using CaCoFold [15]. (iii) The alignment shows covariation that indicates a protein-coding exon instead. If the R-scape output shows no covariation, two additional outcomes are possible: (iv) a plausible but unconfirmed RNA structure if the alignment shows little variation (insufficient power) or (v) an unlikely conserved RNA structure when there is variation (sufficient power) but not covariation. For confirmed conserved RNA structures with sufficient covariation, alignments are refined using Infernal [12].

**Figure 1:**
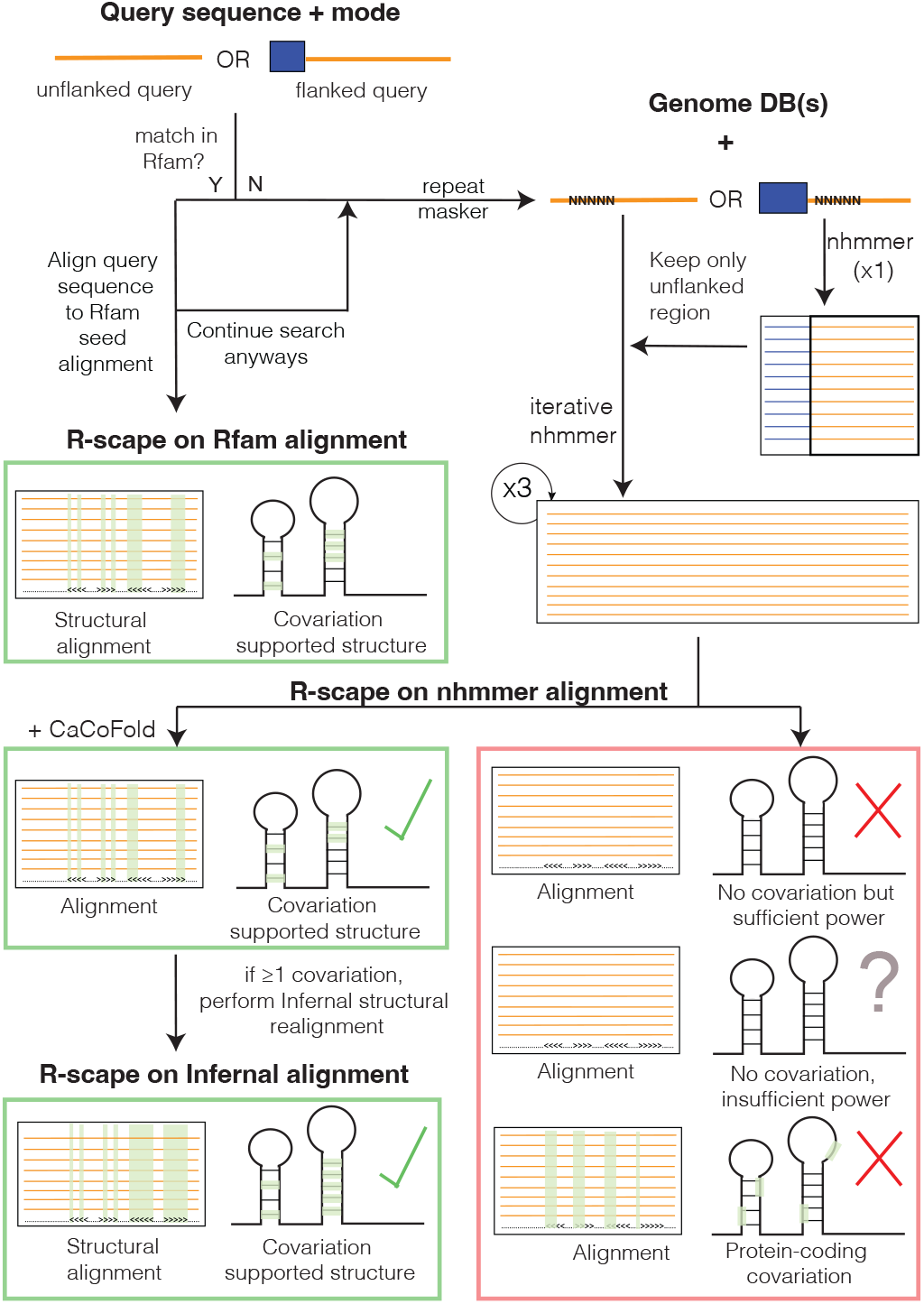
The RNAhub web server workflow for the analysis of an RNA sequence involves assessment of conserved RNA structure by covariation detection. The process starts with a query sequence, either flanked or unflanked. If the query matches an existing RNA family in Rfam, it is aligned directly with the corresponding Rfam seed sequences. Otherwise, the sequence undergoes repeat masking, followed by iterative nhmmer searches against genome databases. The resulting nhmmer alignment is subject to covariation analysis using R-scape to assess covariation support for a conserved RNA structure. In the presence of covariation supports, an structural Infernal alignment is constructed based on the CaCoFold structure.

### Initial search against Rfam

First, the server searches the Rfam database [14]. If there is a hit to the Rfam database, the query sequence and seed sequences from Rfam are used to create a final alignment. In this alignment, the query sequence is merged into an alignment with the Rfam seed sequences according to the Rfam annotated structure (using the Rfam covariance model). R-scape is then executed on this alignment, generating a covariation-optimized structure (see Figure 2a). The user is prompted to decide whether to still proceed with a genome-wide search by clicking Continue the search!. If confirmed, the iterative search is performed (see next section).

**Figure 2:**
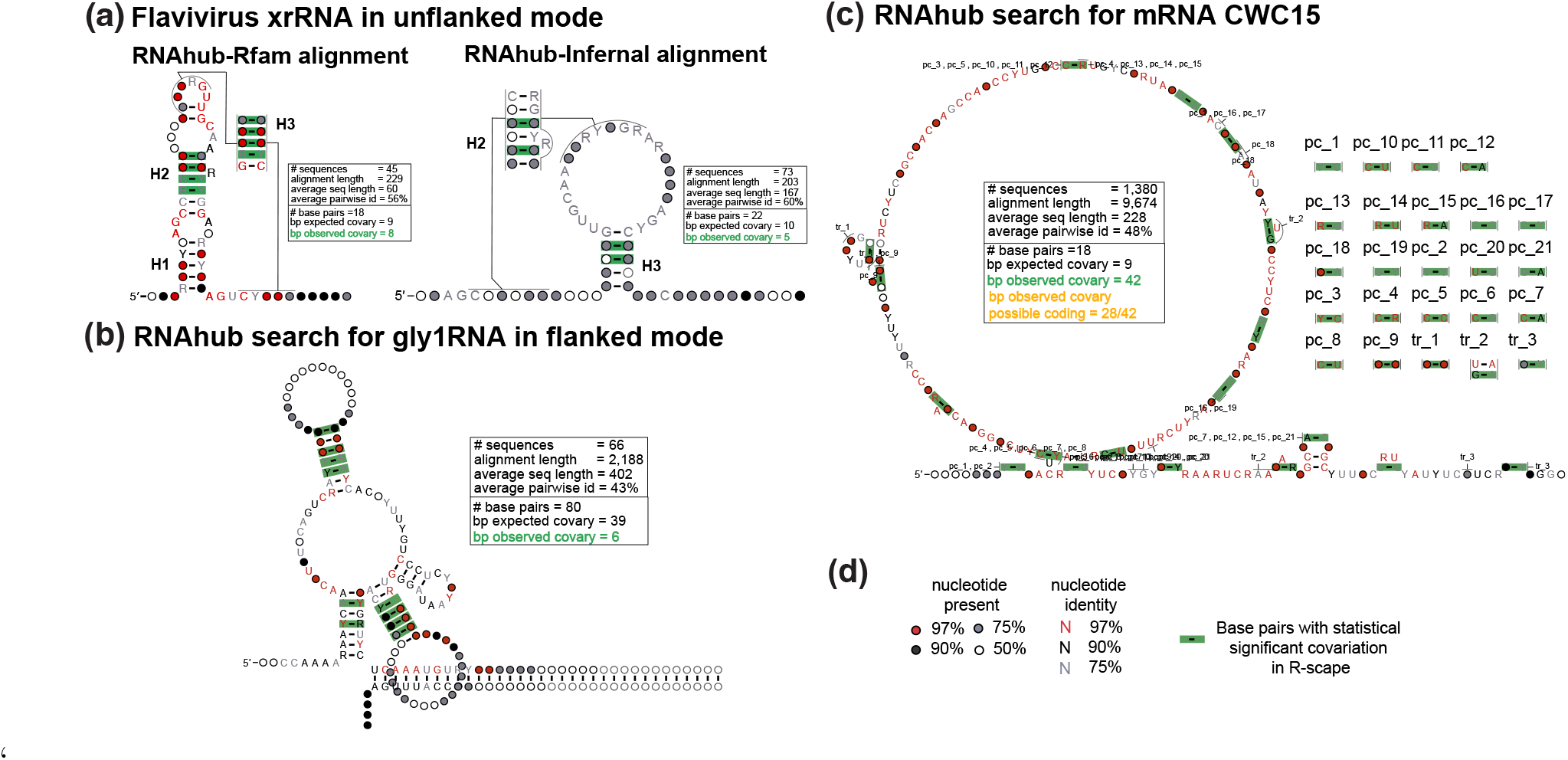
Structural analysis of RNAhub alignments. (a) Structural models of a Flavivirus exoribonuclease-resistant RNA (xrRNA). Comparison of the RNAhub Rfam-based alignment (RF04222) to the RNAhub alignment produced ignoring the known structure searching the viral database with nhmmer and realigning with Infernal. (b) RNAhub structure prediction for the GLY1 RNA motif in the flanked mode with covariation-supported base pairs including three helices. (c) RNAhub results for a protein-coding sequence CWC15. The alignment includes pairs that covary not as a result of RNA base pairing but as a result of protein-coding signal (pc). (d) Legend for nucleotide identity, presence, and base pairs with statistically significant covariation (green rectangles) with R-scape. The RNAhub alignments are provided in the Supplemental Material.

### Iterative search against genome databases

If no match is found in Rfam or the user chooses to continue the genome-wide search, three iterative homology searches are conducted using nhmmer against the database(s) selected in the submitted web form. First, to avoid matching repetitive genomic elements, query sequences are analyzed using RepeatMasker [19] using the Dfam database [24]. Identified repeats are masked with a string of Ns. Then, the first two nhmmer searches are carried out at E-value thresholds of 10^−10^ and the third at 10^−5^. The final alignment obtained after the three iterative homology searches is evaluated using R-scape listing base pairs with covariation support and an R-scape/CaCoFold secondary structure, which integrates all covariation signals into one structural prediction. There are two homology search modes implemented in RNAhub: unflanked and flanked. The unflanked mode uses the given query sequence as input to the iterative homology search. In contrast, the flanked mode first performs a homology search and reports an alignment including flanking regions at one or both ends of the query sequence, which the user must specify. After performing one iteration using the extended (flanking) region, the resulting alignment is trimmed to the input sequence, and the rest of the homology search proceeds as in the unflanked mode. The flanked mode is often advantageous in cases where the query sequence is located within an intron or untranslated region (UTR) of an mRNA or viral RNA. Since the primary sequence of translated coding sequences is typically more conserved at the primary sequence level than introns and UTRs, we recommend using a neighboring exon as a flank if the query sequence is an intron or UTR.

### Covariation analysis and identification of the consensus secondary structure

The server uses R-scape to detect a covariation-supported structure (if any) within the query sequence. As mentioned earlier, the resulting alignments are subjected to the R-scape covariation analysis. R-scape identifies statistically significant covarying base pairs (E-value: 0.05), and it is followed by the method CaCoFold [15], which generates a structural prediction that includes all significantly covarying base pairs, including pseudoknots and other interactions.

R-scape also calculates the power of the alignment to detect covariation which later can be compared to the actual observed covariation [17]. For all base pairs in a structure, we estimate the number of substitutions based on a maximum-likelihood tree [6]. To relate substitutions to power to detect covariation, we used Rfam seed alignments to determine the empirical probability distribution that an Rfam-annotated base pairs with s substitutions shows significant covariation. This yields a curve of covariation power as a function of the number of substitutions. For alignments for which there is a known structure, as it is the case of the RNAhub alignments that show homology to Rfam, we estimate alignment power as the sum of the powers across all possible base pairs. For alignments without a known structure, as it is the case of the RNAhub nhmmer alignments, we estimate power from a structure predicted from the alignment obtained using CaCoFold. Power in the absence of covariation is evidence against the presence of a conserved RNA structure.

### Detection of protein-coding regions

R-scape identifies base pairs that significantly covary above phylogenetic expectation. In addition to structural RNA covariation, another evolutionary signature beyond the phylogenetic null model is protein-coding covariation. These covariations reflect the codon triplet sequence and the evolution of protein-coding sequences in which nucleotides evolve across species in accordance with the maintenance of amino acid identity or functional similarity (see BLOSUM matrix [10]). In contrast to structural RNA covariations, the strongest codon-related covariations are typically within-codon (<3 nucleotides apart). There are within-codon covariations that are >4 nucleotides, similar to structural RNA covariations but differ from structural RNA covariations because they do not conform with canonical base pairing (stem-loops, etc.). See Gao *et al*. 2022 [8] for more information on our characterization of protein-coding covariations. In short, R-scape identifies if a query sequence likely contains a region with protein-coding signatures. For example, R-scape on the nhmmer alignment identifies numerous within-coding covariations, which are denoted as “pc_” in the structure (see Figure 2c).

### Clustering of genomic databases

We provide genome databases for multiple taxa, including eukaryotic clades (metazoa, plants, fungi) and non-eukaryotic clades (bacteria, archaea, viral sequences) downloaded from NCBI [18]. We also provide a metagenomic database obtained from the IMG/VR database [2]. For the smaller non-eukaryotic clades, our databases include all NCBI archaea and viral genomes and all NCBI reference bacterial genomes as of November 2024. To reduce the database size for the larger eukaryotic clades and speed up the search, we subclustered the reference genomes of each clade according to RNase MRP sequence identity and kept one genome per cluster (see Figure S1). These smaller sub-clustered genomes significantly reduced storage and compute time while minimally decreasing the number of covariations for a set of known structural RNAs (U1-U3). Additionally, we provide a structured mammalian genome database to obtain mammal-focused alignments. Additional information about the genome subclustering is provided in the Supplemental Material.

### Implementation

The RNAhub code is composed of two elements: the implementation of the pipeline, rnahub.py, and the web server. rnahub.py was implemented in Python 3, and it is a pipeline that executes nhmmer (3.3.2 at the time of publication) and R-scape (v2.5.6 at the time of publication) and controls the information flow between different steps (Figure 1) and it is based on the reliable code used before for http://rna-tools.online [11]. The web server uses the Django framework for the backend, while the front end uses the jQuery/JavaScript library to send and retrieve the data from the backend. RNAhub.org collects the user inputs within a form and sends the data to the backend (rnahub.py) when the user presses the Search button to start the computation. In the backend, rnahub.py starts on the server command line to perform the search and analysis requested by the user. The output is sent back to the users page, where key results are visualized. Completed result files and a log file are available for download.

## Results

We propose a user-friendly web interface that simplifies the decision-making process for sequence retrieval and alignment. Users have the option to search databases based on phylogenetic origin, as well as to include flanking regions that might be more conserved in the vicinity of the RNA sequence of interest. We also include summaries of alignment statistics, a visual of the predicted RNA structure, and the ability to download alignments for further manipulation.

### Case studies

In the following section, we provide a detailed explanation of RNAhubs functionality by presenting four distinct examples and operational modes. These examples illustrate the platforms capabilities and are visually represented in Figure 2. Details of the analyses with RNAhub outputs are provided in the Supplemental Material.

### xrRNA: search using the unflanked mode

The first example illustrates a full search using the unflanked mode to obtain an alignment and to predict a secondary structure (including the correct prediction of the pseudoknot). The structural exoribonuclease-resistant RNA (xrRNA) corresponds to the intergenic region (IGR) of the Beet western yellows virus (BWYV) genomic sequence (GeneBank accession no. NC_004756), which was earlier reported to contain the xrRNA structure by Steckelberg *et al*. [21]. This structural RNA has already been incorporated as a Rfam family (RF04222). As indicated by the RNAhub output, users have the option to either continue with the standard

RNAhub workflow or directly align the query sequence using the corresponding Rfam seed alignment. When analyzing the Rfam alignment with R-scape (Figure 2a), the proposed secondary structure comprises 18 base pairs, with 8 observed to covary (expected covariation: 8.7). The user can also ask for the interactive search to generate alignment from scratch. The Infernal alignment analyzed by R-scape proposes a larger structure of 37 base pairs but identifies only 4 covarying pairs (from 20 expected to covary). These results highlight that the Rfam-based alignments provide more accurate and reliable predictions of RNA secondary structures. However, in the absence of that knowledge, the RNAhub nhmmer search against the viral genomes reports an alignment that reflects well the structural constraints of the molecule.

### GLY1 RNA: search using the flanked mode

The second example illustrates the use of RNAhub’s flanked mode to enhance homology searches, enabling the detection of structural RNAs that would otherwise remain undetected. Here, the RNAhub flanked mode is demonstrated through analysis of the the 3’ UTR of the GLY1 gene from *Saccharomyces cerevisiae*, replicating the results reported by Gao *et al*. 2021 [7]. The flanked mode is particularly useful for identifying homologous sequences of rapidly evolving structural RNAs, which might be difficult to detect directly due to low primary sequence conservation. By including adjacent, more conserved regions (flanking regions) in the search, RNAhub enhances alignment quality and facilitates the discovery of conserved RNA structures. This mode is especially beneficial for analyzing introns and untranslated regions (UTRs) of mRNAs, where many structural RNAs, such as riboswitches, perform essential regulatory functions. The query sequence, along with its flanking region, corresponds to a segment of chromosome V (chrV), spanning positions 61,799 to 73,793 (11.99 kb). The whole region encompasses the GLY1 gene and its associated untranslated region (UTR). GLY1 encodes threonine aldolase, an enzyme responsible for catalyzing reactions involving threonine metabolism. The R-scape analysis of the Infernal alignment proposes a secondary structure comprising 59 base pairs, of which 10 are observed to covary (8.7 covarying pairs are expected) (Figure 2b). These results emphasize that the use of flanking regions is crucial for identifying conserved structural RNAs, as this RNA structure would otherwise remain undetected due to rapid sequence divergence.

### CWC15: protein-coding covariation detected

We conducted an unflanked mode search using a protein-coding query sequence of CWC15, one of the spliceosomal proteins found in yeast, to demonstrate how R-scape can also detect protein-coding covariations. As expected, no hits were found against Rfam, confirming that the query is not a structural RNA. Instead, the covariation signal is most consistent with a protein-coding region rather than a structural RNA (Figure 2c). The presence of statistically significant covariation suggests a protein-coding sequence. The proposed secondary structure contains zero base pairs, while 42 base pairs exhibit covariation (because of protein-coding covariations).

### HOTAIR: a long non-coding RNA with evolutionary evidence against a conserved structure

We used the long non-coding RNA HOTAIR as an example of an RNA conserved in mammalian species, where the alignment has enough power but not covariation that can be associated with a conserved RNA structure. We conducted a homology search for domain 1 of HOTAIR using a database of 94 mammalian genomes. The final alignment created by the RNAhub has sequences from 92 genomes with an average percentage identity of 66%. The R-scape analysis of this alignment indicates that while the alignment expects 61 base pairs to covary, only 7 are found, and their arrangement does not support any RNA helix. Thus, the RNAhub alignment for HOTAIR-D1 (as others before [16, 17]) presents evidence against HOTAIR displaying a conserved RNA structure.

## Discussion

The RNAhub web server streamlines the process of obtaining accurate RNA structural models by integrating evolutionary information through a user-friendly automated workflow. Unlike other commonly employed approaches, RNAhub does not necessarily produce an output structure for every RNA sequence submitted. Instead, the absence of a predicted structure reliably indicates insufficient evolutionary variability or inadequate covariation support rather than a limitation of the computational pipeline itself. This selective approach differentiates RNAhub from methods that invariably output secondary structure, regardless of confidence. RNAhub typically delivers a single consensus RNA structure. Importantly, even if the predicted secondary structures may not always precisely represent the final three-dimensional RNA fold, the covariation-supported structural information identified by RNAhub remains valuable. Such evolutionary insights can significantly enhance the predictive accuracy of complementary computational modeling approaches, thereby facilitating more targeted and precise downstream experimental designs. In conclusion, RNAhub provides a robust, automated solution for identifying structurally conserved RNAs based on rigorous evolutionary analyses. Its precise and selective methodology significantly advances RNA research by reducing uncertainty and enhancing confidence in structural predictions, reinforcing its role as a critical tool for both computational and experimental RNA studies.

## Data availability

The web server is available at https://rnahub.org. This website is free and open to all users without a login requirement.

## Authors contribution

**Marcin Magnus**: Conceptualization, Methodology, Software, Writing - Original Draft, Writing - Review & Editing. **William Gao**: Conceptualization, Methodology, Software, Writing - Original Draft,Writing - Review & Editing. **Nivedita Dutta**: Validation, Writing - Original Draft,Writing - Review & Editing. **Quentin Vicens**: Conceptualization, Supervision, Writing - Original Draft, Writing - Review & Editing. **Elena Rivas**: Conceptualization, Supervision, Writing - Original Draft, Writing - Review & Editing.

## Acknowledgments

We would like to thank Jeff Kieft for early discussions and support, Ann Yang for early contributions to this work, and Agata Kilar for support in the development of this manuscript.

---

**Figure S1:**
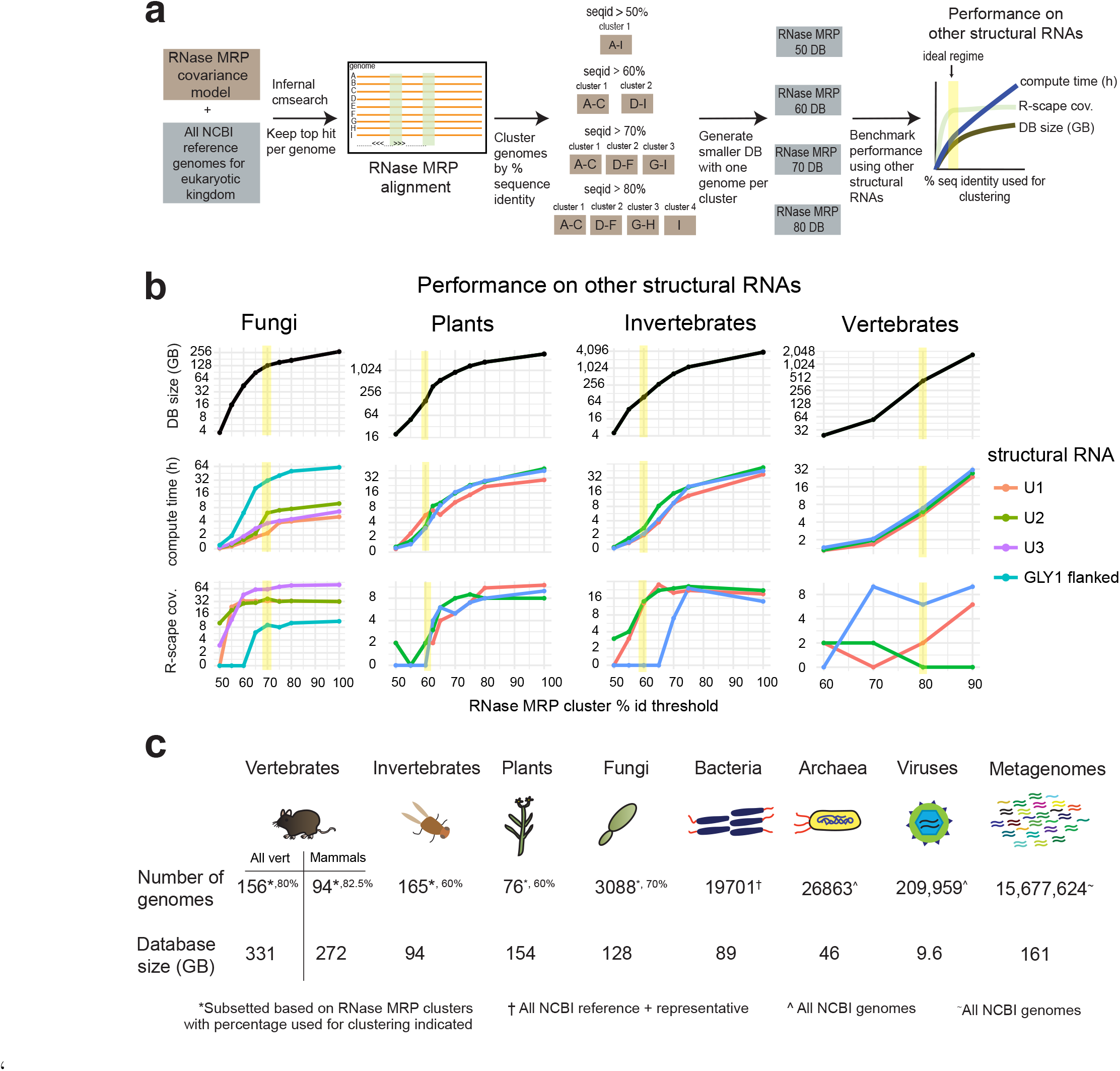
Clustering of genomes. (a) To reduce the size of eukaryotic databases, we clustered genomes based on their RNase MRP sequence (a highly conserved structural RNA), and kept one per cluster to construct smaller genome databases. We determined the ideal regime as the subclustered genome database that maximizes the tradeoff between the number of R-scape covariations vs. compute time and database (DB) size. (b) Database size, compute time, and R-scape covariation tradeoff (in log scale) for three non-RNase MRP structural RNAs using clustering thresholds ranging from 0.50 to 1.00, in increments of 0.05. The ideal regime that balances covariation against DB size and compute time is highlighted in yellow (c) Genome databases available in RNAhub, based on either the clustering at the ideal regime for eukaryotic genomes, or all reference NCBI for bacteria, and all NCBI archaea and viral genomes.

