## Supplementary material for "RNAhub - an automated pipeline to search and align RNA homologs with secondary structure assessment": Genome_subclustering_FigureS1b.pptx

### Slide 1
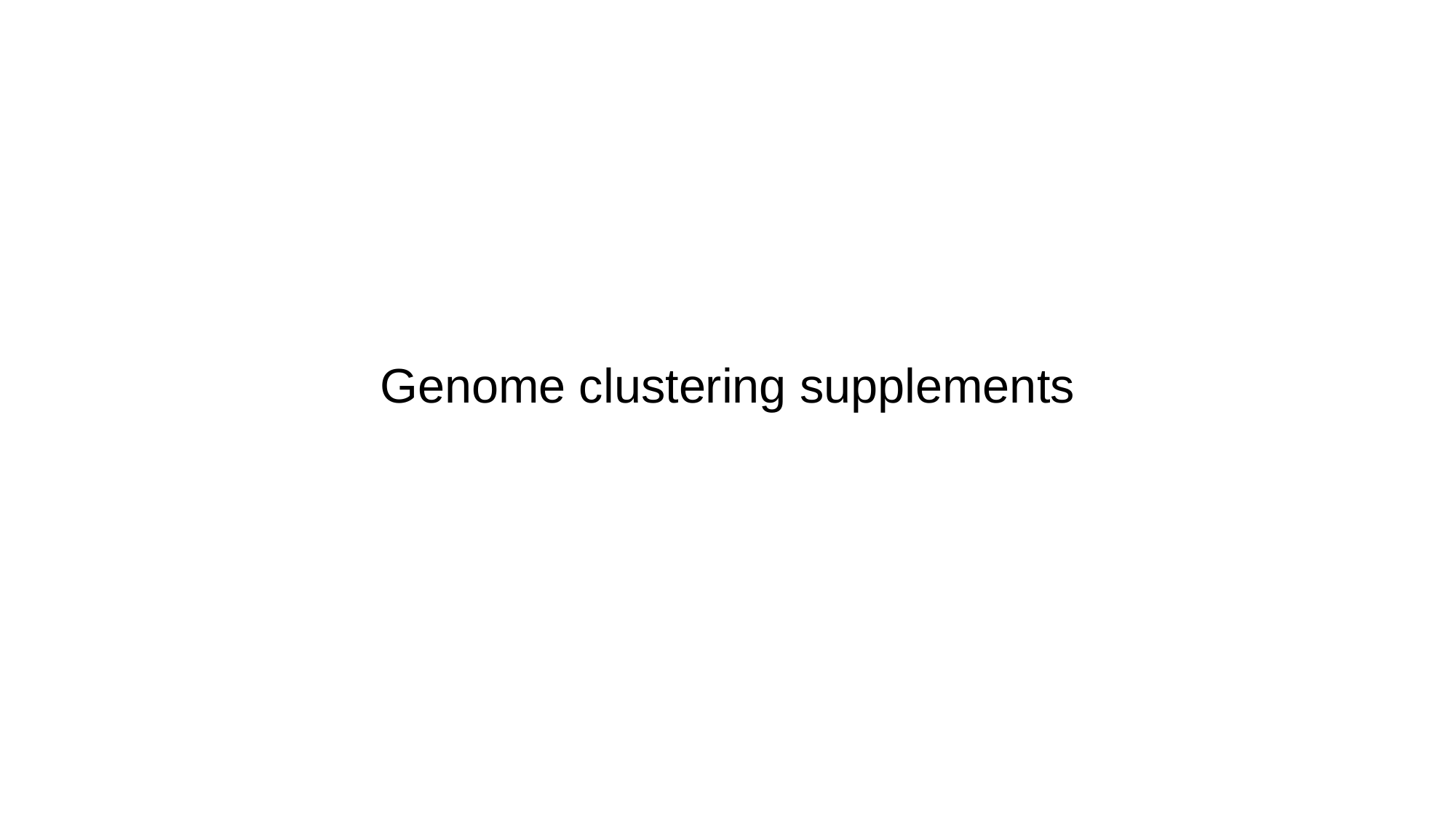

#### Genome clustering supplements

### Slide 2
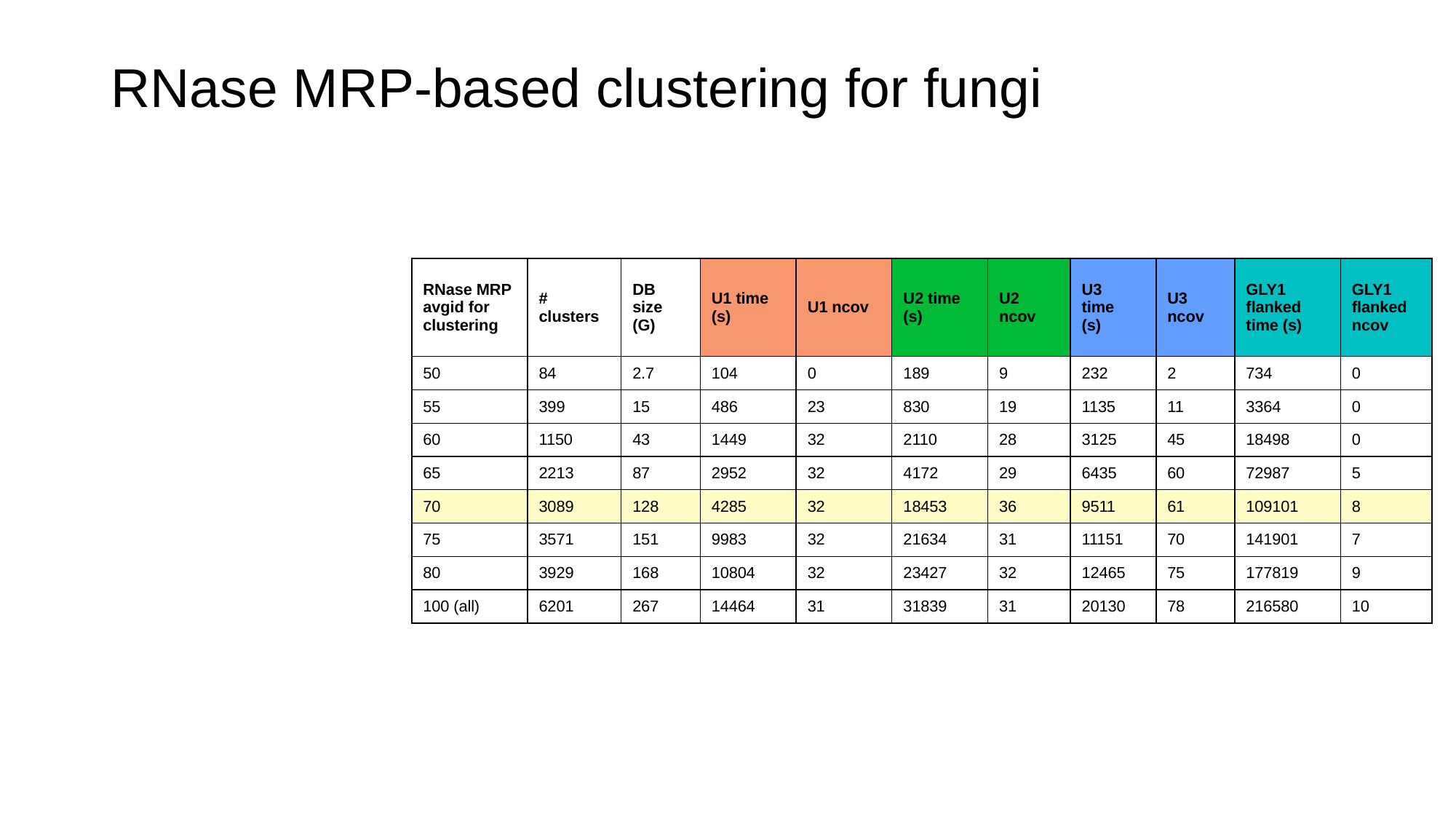

#### RNase MRP-based clustering for fungi
| RNase MRP avgid for clustering | # clusters | DB size (G) | U1 time (s) | U1 ncov | U2 time (s) | U2 ncov | U3 time (s) | U3 ncov | GLY1 flanked time (s) | GLY1 flanked ncov |
| --- | --- | --- | --- | --- | --- | --- | --- | --- | --- | --- |
| 50 | 84 | 2.7 | 104 | 0 | 189 | 9 | 232 | 2 | 734 | 0 |
| 55 | 399 | 15 | 486 | 23 | 830 | 19 | 1135 | 11 | 3364 | 0 |
| 60 | 1150 | 43 | 1449 | 32 | 2110 | 28 | 3125 | 45 | 18498 | 0 |
| 65 | 2213 | 87 | 2952 | 32 | 4172 | 29 | 6435 | 60 | 72987 | 5 |
| 70 | 3089 | 128 | 4285 | 32 | 18453 | 36 | 9511 | 61 | 109101 | 8 |
| 75 | 3571 | 151 | 9983 | 32 | 21634 | 31 | 11151 | 70 | 141901 | 7 |
| 80 | 3929 | 168 | 10804 | 32 | 23427 | 32 | 12465 | 75 | 177819 | 9 |
| 100 (all) | 6201 | 267 | 14464 | 31 | 31839 | 31 | 20130 | 78 | 216580 | 10 |

### Slide 3
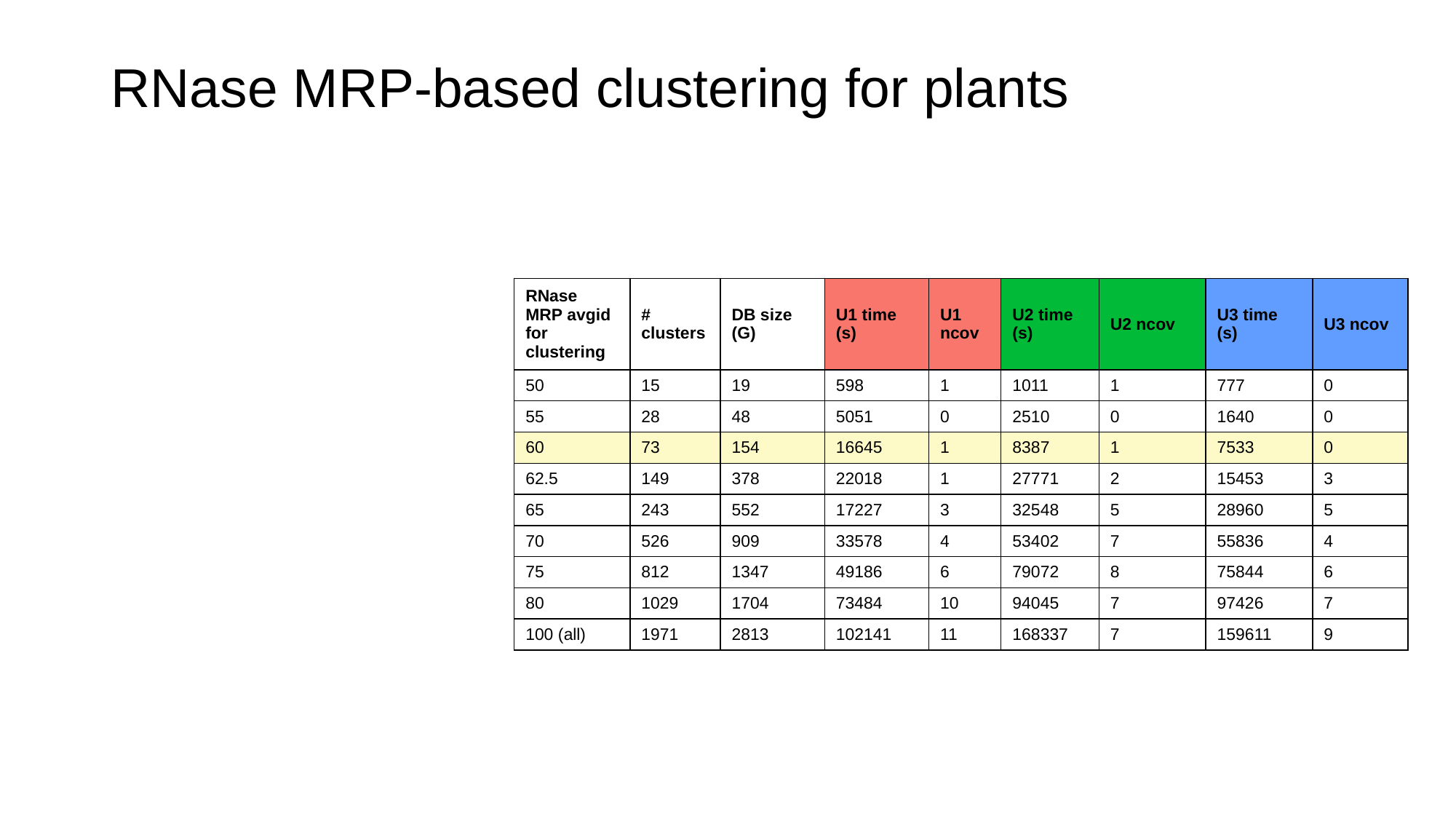

#### RNase MRP-based clustering for plants
| RNase MRP avgid for clustering | # clusters | DB size (G) | U1 time (s) | U1 ncov | U2 time (s) | U2 ncov | U3 time (s) | U3 ncov |
| --- | --- | --- | --- | --- | --- | --- | --- | --- |
| 50 | 15 | 19 | 598 | 1 | 1011 | 1 | 777 | 0 |
| 55 | 28 | 48 | 5051 | 0 | 2510 | 0 | 1640 | 0 |
| 60 | 73 | 154 | 16645 | 1 | 8387 | 1 | 7533 | 0 |
| 62.5 | 149 | 378 | 22018 | 1 | 27771 | 2 | 15453 | 3 |
| 65 | 243 | 552 | 17227 | 3 | 32548 | 5 | 28960 | 5 |
| 70 | 526 | 909 | 33578 | 4 | 53402 | 7 | 55836 | 4 |
| 75 | 812 | 1347 | 49186 | 6 | 79072 | 8 | 75844 | 6 |
| 80 | 1029 | 1704 | 73484 | 10 | 94045 | 7 | 97426 | 7 |
| 100 (all) | 1971 | 2813 | 102141 | 11 | 168337 | 7 | 159611 | 9 |

### Slide 4
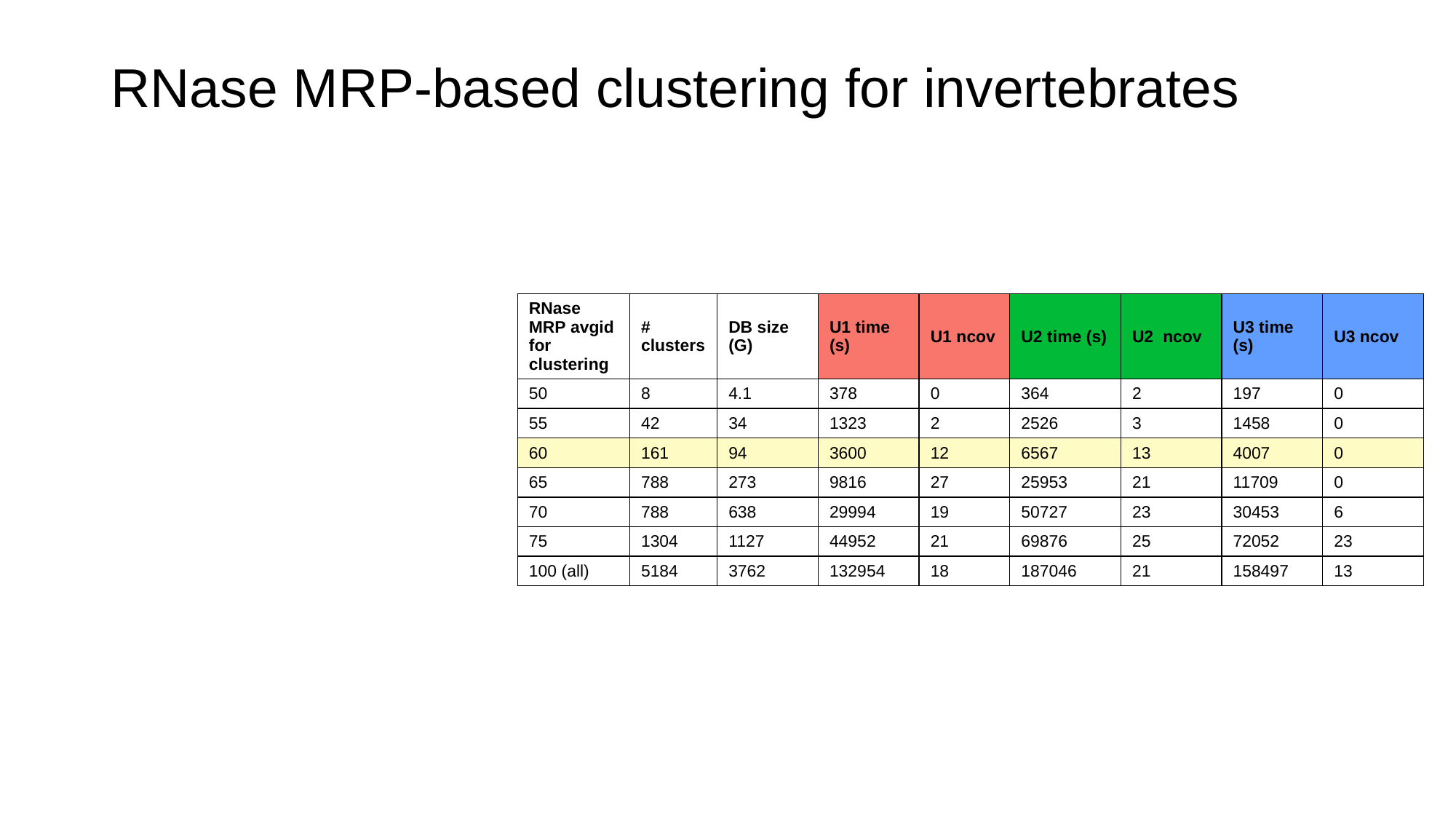

#### RNase MRP-based clustering for invertebrates
| RNase MRP avgid for clustering | # clusters | DB size (G) | U1 time (s) | U1 ncov | U2 time (s) | U2 ncov | U3 time (s) | U3 ncov |
| --- | --- | --- | --- | --- | --- | --- | --- | --- |
| 50 | 8 | 4.1 | 378 | 0 | 364 | 2 | 197 | 0 |
| 55 | 42 | 34 | 1323 | 2 | 2526 | 3 | 1458 | 0 |
| 60 | 161 | 94 | 3600 | 12 | 6567 | 13 | 4007 | 0 |
| 65 | 788 | 273 | 9816 | 27 | 25953 | 21 | 11709 | 0 |
| 70 | 788 | 638 | 29994 | 19 | 50727 | 23 | 30453 | 6 |
| 75 | 1304 | 1127 | 44952 | 21 | 69876 | 25 | 72052 | 23 |
| 100 (all) | 5184 | 3762 | 132954 | 18 | 187046 | 21 | 158497 | 13 |

### Slide 5
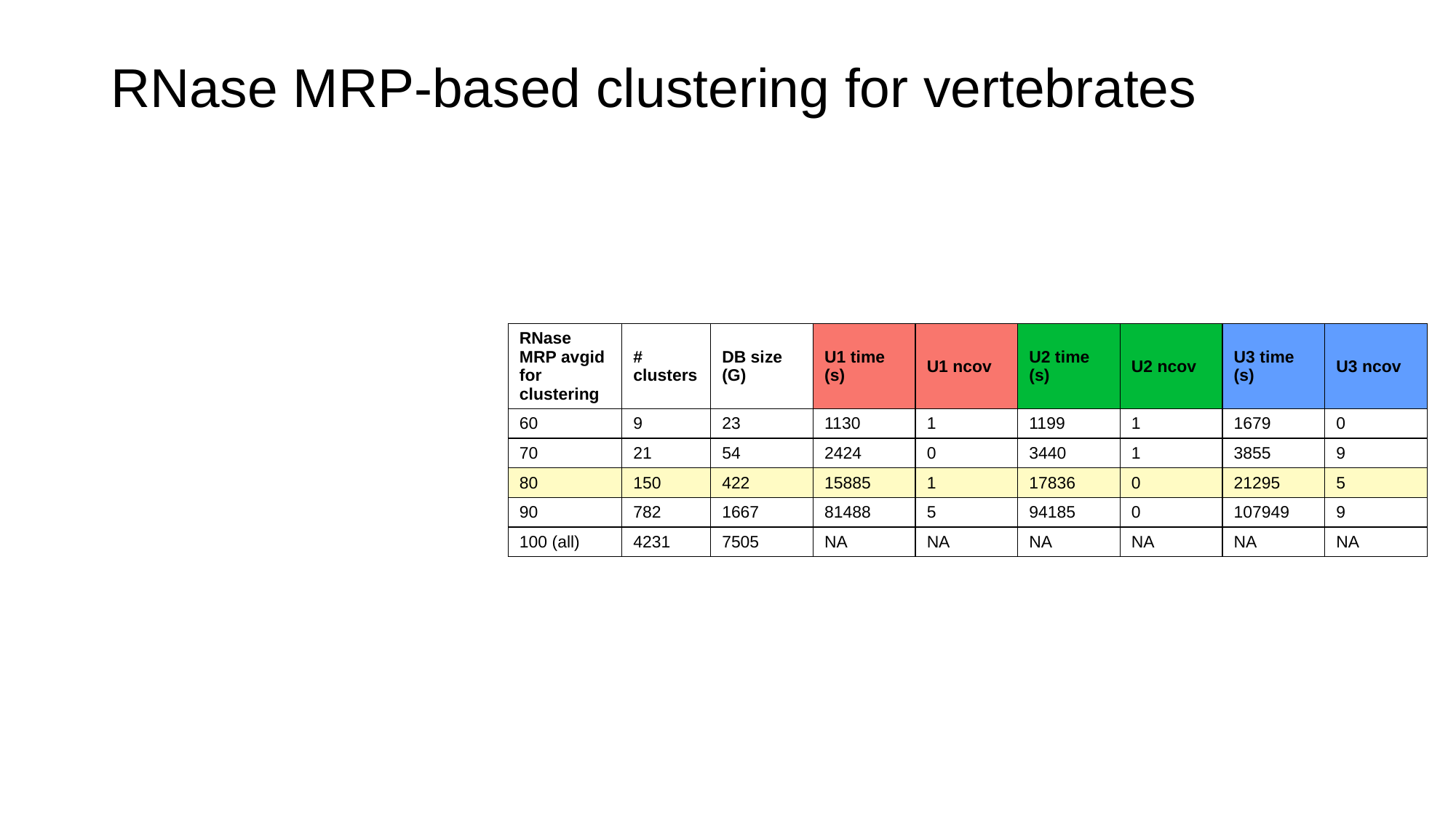

#### RNase MRP-based clustering for vertebrates
| RNase MRP avgid for clustering | # clusters | DB size (G) | U1 time (s) | U1 ncov | U2 time (s) | U2 ncov | U3 time (s) | U3 ncov |
| --- | --- | --- | --- | --- | --- | --- | --- | --- |
| 60 | 9 | 23 | 1130 | 1 | 1199 | 1 | 1679 | 0 |
| 70 | 21 | 54 | 2424 | 0 | 3440 | 1 | 3855 | 9 |
| 80 | 150 | 422 | 15885 | 1 | 17836 | 0 | 21295 | 5 |
| 90 | 782 | 1667 | 81488 | 5 | 94185 | 0 | 107949 | 9 |
| 100 (all) | 4231 | 7505 | NA | NA | NA | NA | NA | NA |
