## Supplementary figures and images for "RNAhub - an automated pipeline to search and align RNA homologs with secondary structure assessment"

### CWC15.cacofold.R2R.sto.pdf

CWC15.cacofold

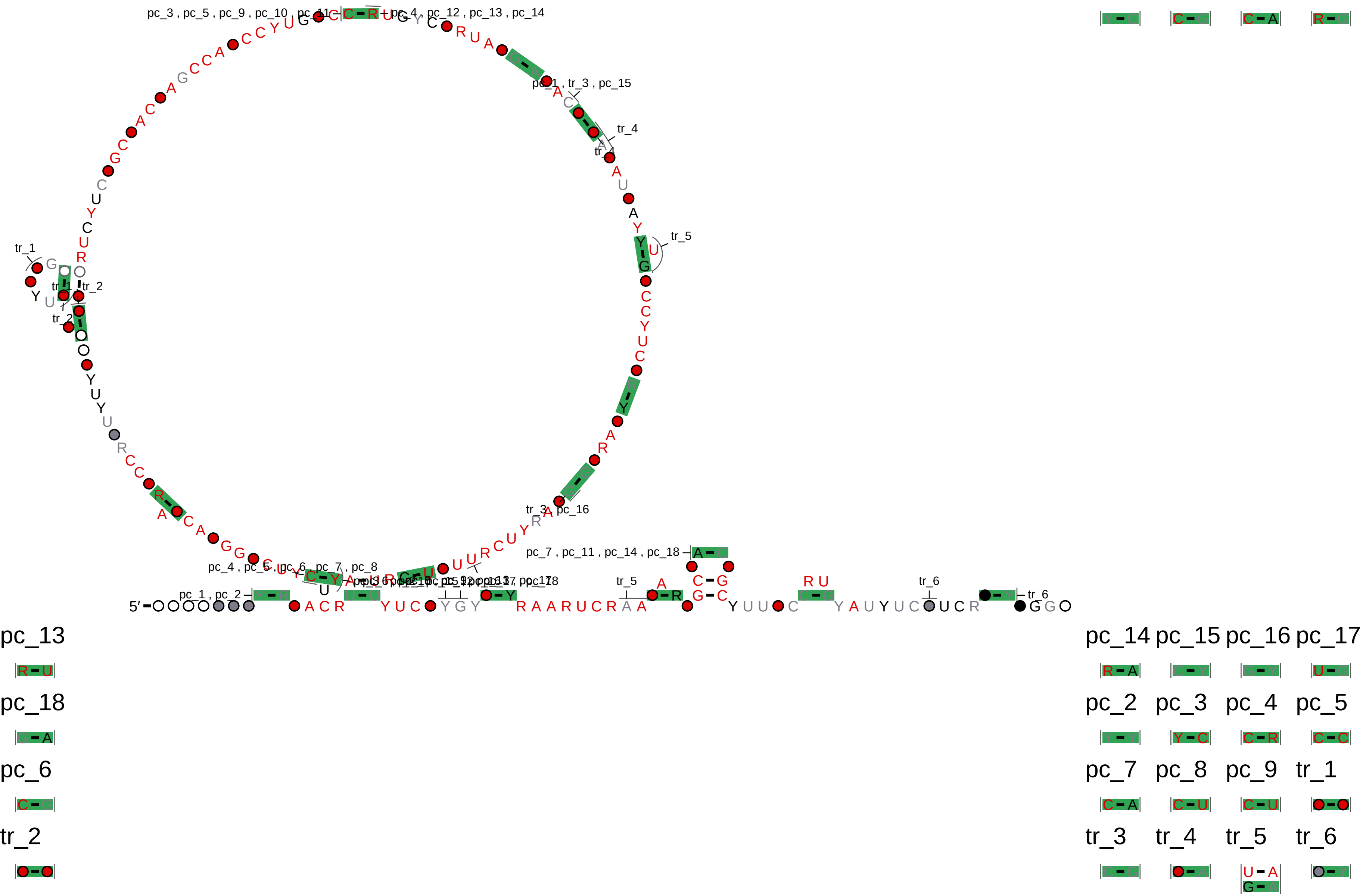

### GLY.cacofold.R2R.sto.pdf

GLY.cacofold

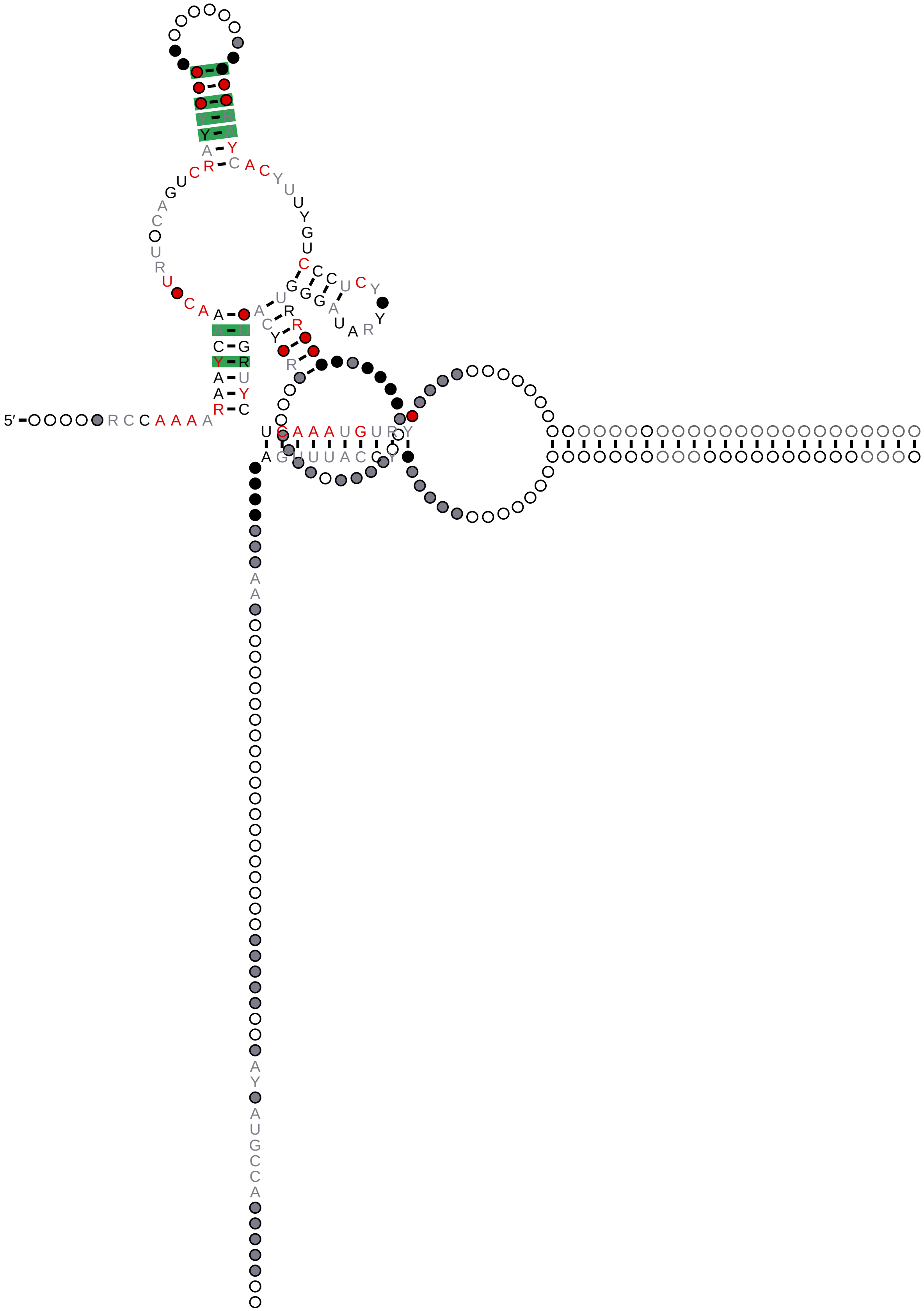

### GLY.R2R.sto.pdf

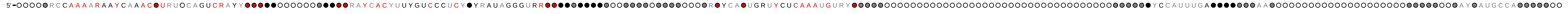

### HOTAIR_D1_Homo_sapiens_1-526.cacofold.R2R.sto.pdf

HOTAIR_D1_Homo_sapiens_1-526.cacofold

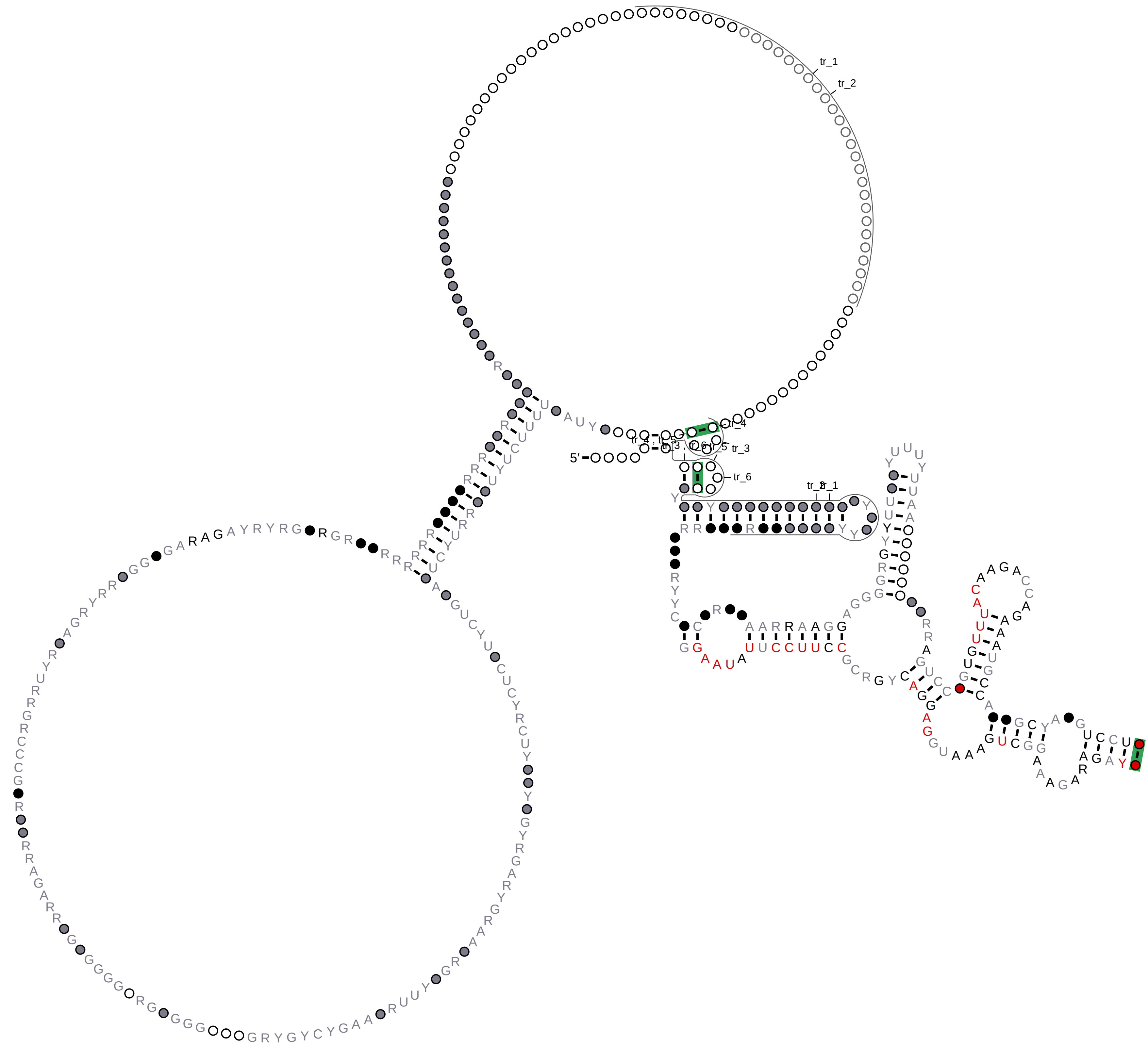


tr_1

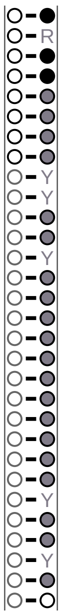

tr_2

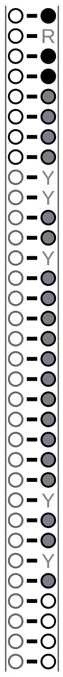

tr_3

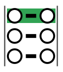

tr_4

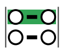

tr_5

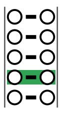

tr_6

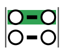

### HOTAIR_D1_Homo_sapiens_1-526.R2R.sto.pdf

#### HOTAIR_D1_Homo_sapiens_1-526

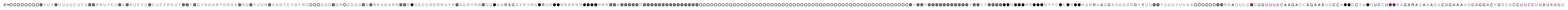

### infernal_1.cacofold.R2R.sto.pdf

#### infernal_1.cacofold

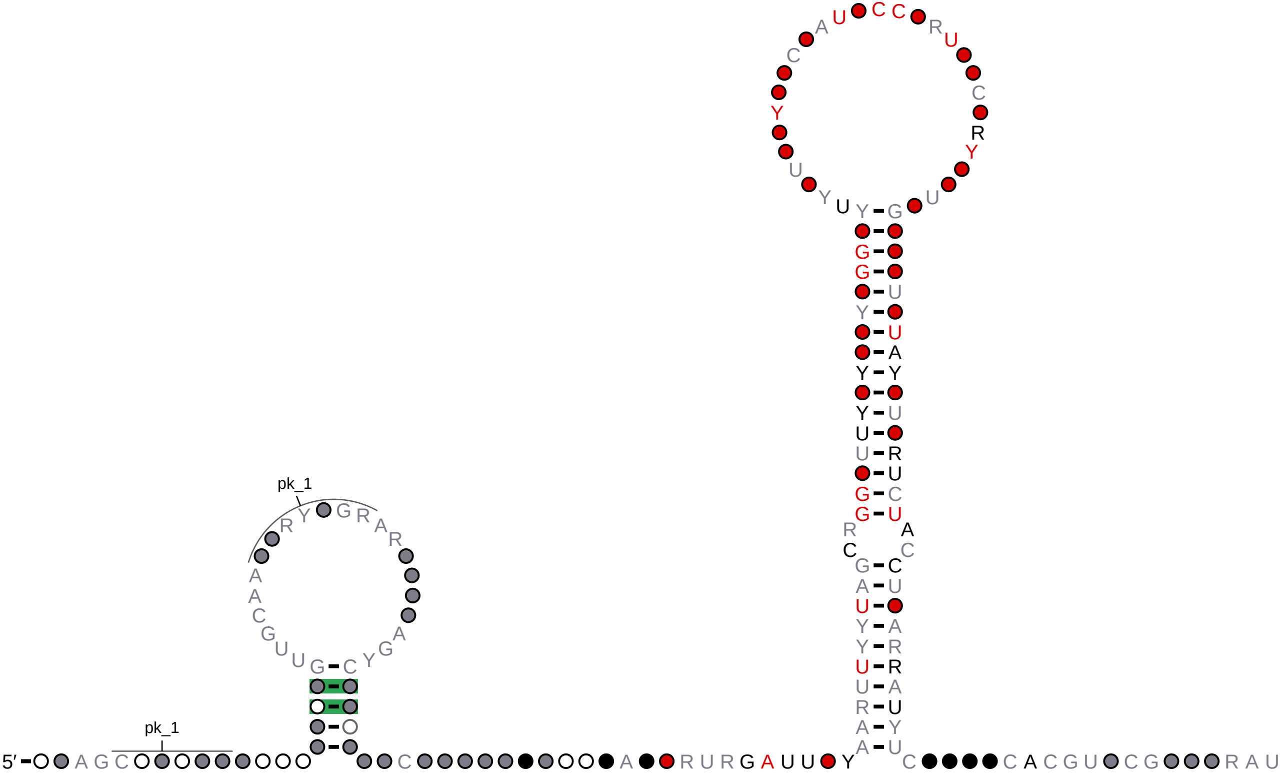


# pk_1

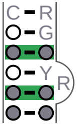

### infernal_1.cacofold.R2R.sto.pdf

infernal_1.cacofold

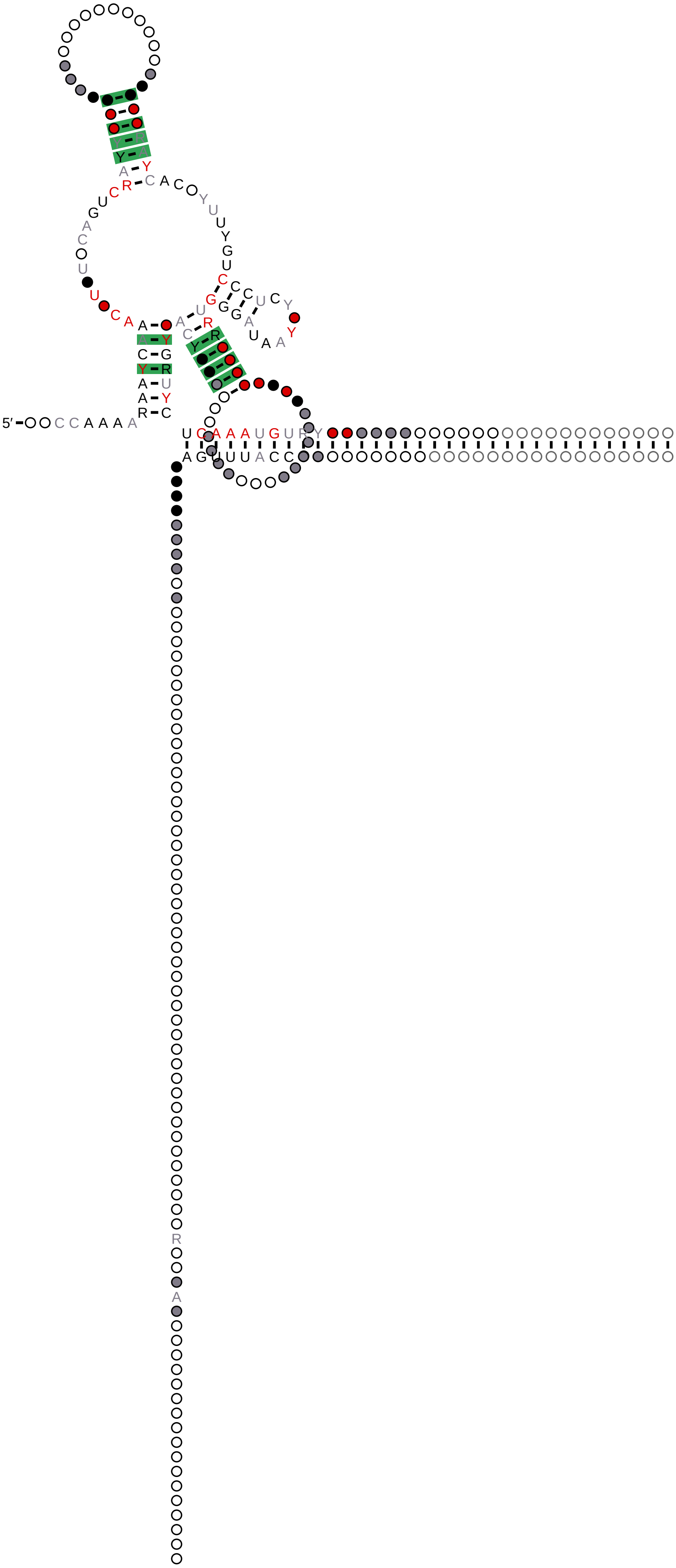

### infernal_1.R2R.sto.pdf

#### infernal_1

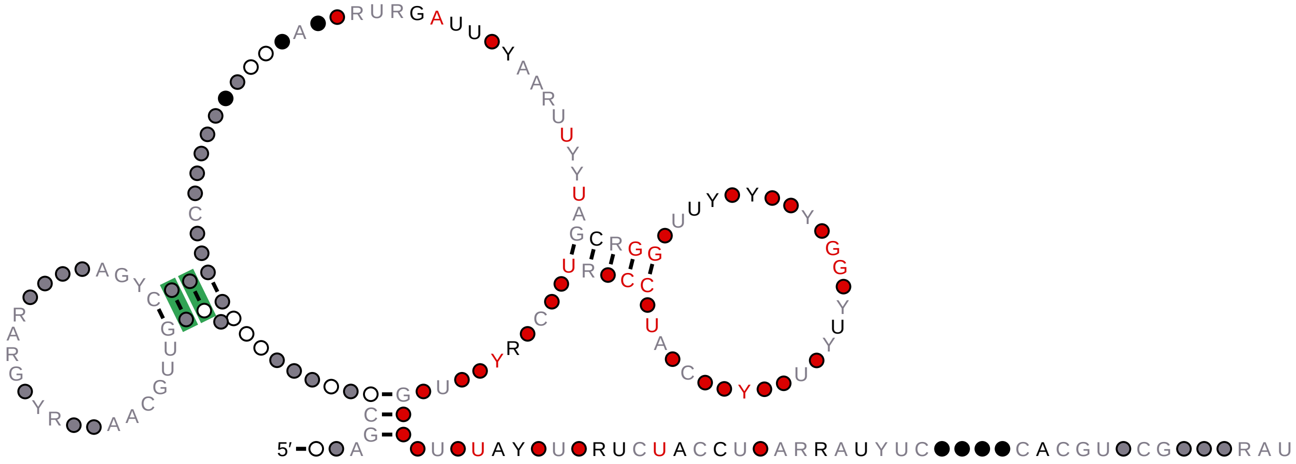

### infernal_1.R2R.sto.pdf

infernal_1

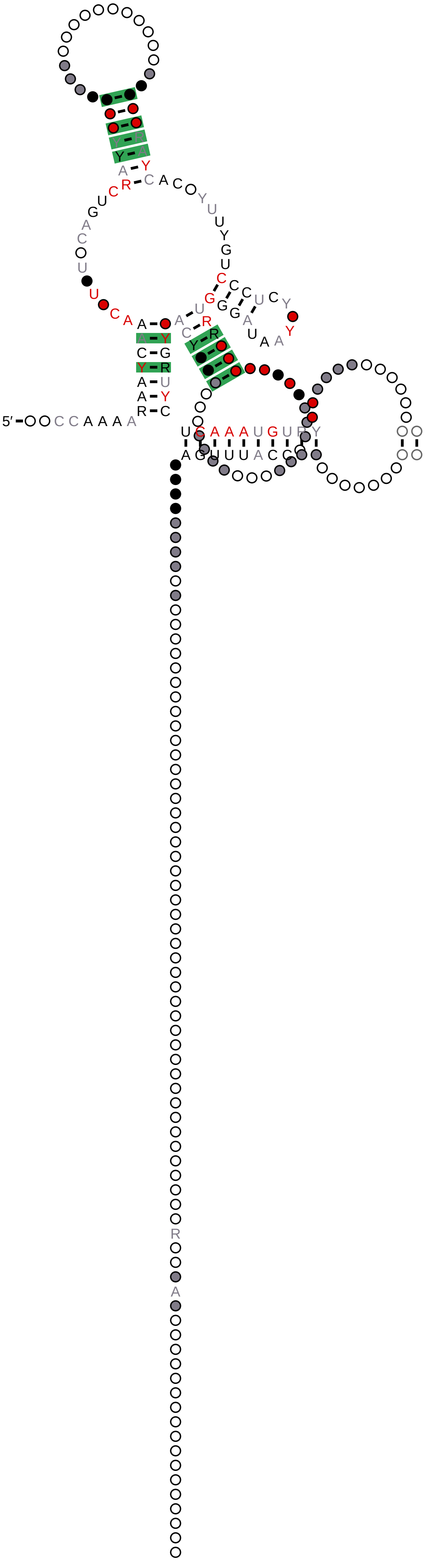

### seq+RF04222_seed_1.cacofold.R2R.sto.pdf

#### seq+RF04222_seed_1.cacofold

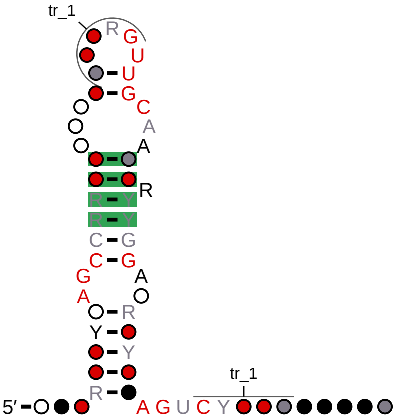


## tr_1

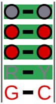

### seq+RF04222_seed_1.R2R.sto.pdf

#### seq+RF04222_seed_1

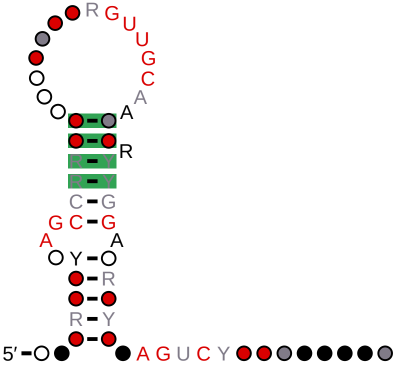

### xrRNA.cacofold.R2R.sto.pdf

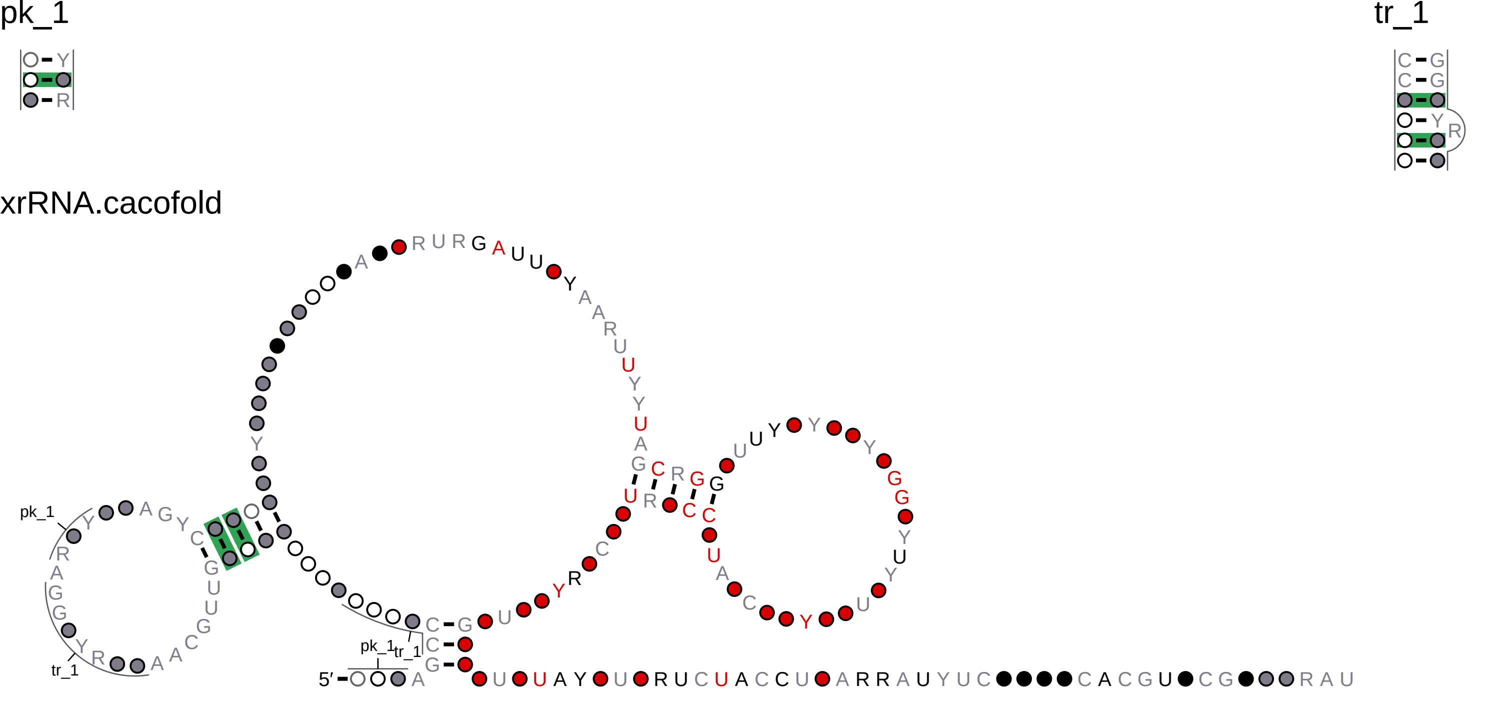

### xrRNA.R2R.sto.pdf

xrRNA

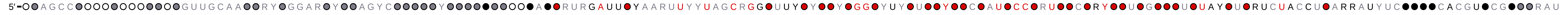
